# taxutils: Tools for managing and operating on NCBI taxonomy and accession mappings

**DOI:** 10.64898/2026.09.18.752821

**Authors:** William O’Brien, Seungmo Lee, Vivek Agarwal, Eleazar Eskin

## Abstract

Research in bioinformatics demands that many workflows and operations be consistently repeated across different tools. These operations include managing a taxdump, dealing with accession mappings, and performing analysis on the taxonomic tree. We introduce `taxutils` to address and alleviate the strain of these common workflows. To exhibit the extent of the package’s capabilities, we introduce a strategy for assessing quality of an assembly via its potential impact on and perturbation of its mapping to a lowest-common-ancestor index. We illustrate the steps required for such an analysis, and show how the tool makes data management easy and otherwise-complex workflows simple.

## 1. Introduction

The NCBI taxonomy is so widely used across bioinformatic tools that a package for its routine operations emerges naturally from the tasks one repeats when working with it. Reference genomes are identified by accession IDs, which are buried within non-standardized header strings. Taxonomic IDs index a hierarchical tree whose nodes are connected by nested relationships, where each node is assigned a rank reflecting the phenotypic traits and average nucleotide index (ANI) threshold that distinguish it from relatives (Ciufo et al., 2018). Working with this data requires parsing accession IDs out of headers, mapping IDs to taxa, filtering by certain properties, and reassembling them into formatted headers.

Tools such as `Kraken2` (Wood et al., 2019), `Bracken` (Lu et al., 2017), `KrakenUniq` (Breitwieser et al., 2018), `Centrifuge` (Kim et al., 2016), `KMCP` (Shen et al., 2023), `ganon` (Piro et al., 2020) each require, prior to database construction, a mapping from sequence or accession identifiers to taxonomic IDs. Abundance estimators that operate over bare sequence headers, such as `Sylph` (Shaw and Yu, 2025), defer taxonomy to the user and require custom post-processing to translate genome accessions into lineages. `Kraken2` expects headers with taxo-nomic ID delimited by `kraken:taxid|` in each sequence header, `Centrifuge` and `KrakenUniq` consume a standalone `seqid2taxid` conversion table, and `KMCP` and `Metabuli` use the assembly accession recovered from the file name or the first header field (Kim and Steinegger, 2024). These accessions are frequently embedded in heterogeneous, inconsistently formatted headers which can lead to unpredictable mapping errors during database construction (jlrolando, 2020; yifengyuan, 2019). Though some amount of the groundwork is automated by these tools, users are left to diagnose and repair these mappings by hand when automation fails (ghbore, 2018; AzizNasr, 2023). Parsing, recovering, and re-emitting these identifiers is therefore recurring, unavoidable groundwork that requires a reliable solution.

`TaxonKit` (Shen and Ren, 2021), `taxopy` (Camargo et al., 2025), and the `NCBITaxa` module of ETE3 (Huerta-Cepas et al., 2016), offer elements of lineage retrieval, rank fetching, and lowest-common-ancestor (LCA) computation. Other tools such as `seqkit2` (Shen et al., 2024) provide general-purpose manipulation of FASTA/FASTQ records and sequences. However, for deep analysis of the taxonomic tree and the accessions which comprise its taxa, these tools lack the flexibility we require: parsing of accession IDs, calculation of tree topological features, taxa selection, rank corrections, and accessible data objects.

To address these issues, we present `taxutils`, a Python framework with a Rust backend that simplifies the management and manipulation of taxonomic data. From the command-line, we offer five core commands: `clean` FASTA headers in-place, `deduplicate` accessions in FASTA records, `extract` accession IDs from FASTA headers, `filter` FASTA records by taxonomic ID, and `grep` FASTA records matching targeted accessions. The rest of the package is designed for users to compose base functions into customized scripts to fit the constantly evolving landscape of bioinformatics tools and data structures. We demonstrate the tool’s flexibility through display of a quality scoring method for reference genomes. We show how `taxutils` can be used to easily evaluate how an assembly perturbs the existing taxonomic hierarchy as a heuristic for assembly quality and contamination. Carrying out this assessment surfaces a range of shortcomings in the RefSeq database and NCBI taxonomy, including inconsistencies in the ranking system, problematic topologies within the tree, and irregularities in the clade structure of different taxonomic subtrees; `taxutils` offers corrections and helps navigate these issues.

We use the tool to present an analysis of GenBank nucleotide viral sequences, identifying families that are underrepresented in RefSeq and producing an approach for curating viruses to fill these gaps in k-mer databases. We additionally show how accession to taxon mappings can be relabeled within our framework of sequence curation.

## 2. Results

### 2.1. Rank corrections address viral family incongruities

The RefSeq viral curation lacks many taxa in the NCBI taxonomic tree that are necessary for high-recall metagenomic classification. Some viral families in the RefSeq database are represented more heavily than others, leading to gaps in the neglected family’s visibility during classification. Identifying these families in the first place can be challenging, due to inconsistencies in the assigned taxonomic ranks. There are instances, for example, where taxa are assigned without rank, parents and children are assigned identical rank, or taxa are assigned outlier rank codes, and so we cannot consistently establish a taxon’s place on the tree relative to its children and ancestors. Fig. 1a illustrates that 9.3% of taxa are assigned no rank, with a further 3.93% assigned to some non-standard category.

**Fig. 1.**
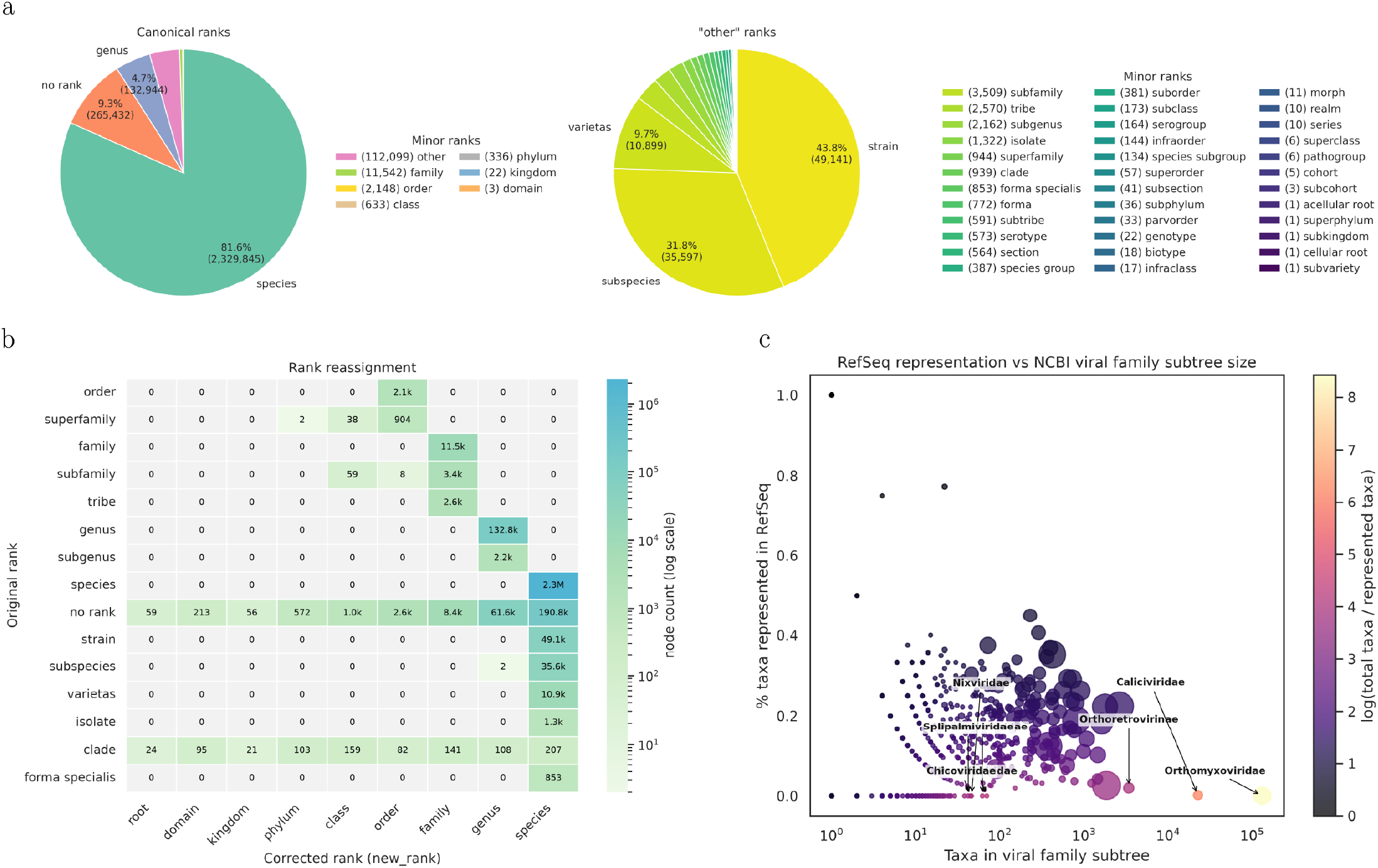
(**a**) On the left, the distribution of the canonical ranks. A significant portion of nodes are assigned “no rank” or fall into a noncanonical rank (“other”). On the right, the distribution of non-canonical ranks. (**b**) Movements of rank assignments for the top 15 largest ranks, after the `taxutils` correction. (**c**) Each point represents a viral family, positioned by its cardinality (x-axis, log scale) against the fraction of its taxa represented in RefSeq (y-axis). Hue encodes the log-ratio of total to represented taxa; large families falling to the lower-right are substantially underrepresented despite their size.

To fix this issue, `taxutils` offers a correction that uses a canonical set of ranks as anchors and assigns the remaining taxa positions from their parent and child anchors (Methods 4.1), following the precedent set in Kraken2 (Wood et al., 2019). Fig. 1b illustrates the reassignments across `taxdump.dmp` for the most prevalent movements. The majority of ambiguous ranks are reassigned to the species level. Crucially, we do not see any canonical ranks reassigned to a different canonical rank, preserving the intended anchors.

### 2.2. Topologies of viral family clades

With properly established ranks, we can find the closest instance of a family viral clade and use this information to analyze the distribution of existing sequences in the RefSeq database across different viral families. Indeed, there are large viral families which are significantly lacking representation in the RefSeq database (Fig. 1c; see full statistics in Sup. S1). Orthomyxoviridae, for instance, contains 131,933 total taxa, with only 28 of its taxa represented in RefSeq (backed by 191 accessions), including a large number of influenza A and B sub-types. The mutation rate of the influenza viruses suggests the divergence of influenza subtypes represented in the taxonomy outpaces our ability to curate clean accessions for RefSeq, necessitating algorithmic supplements for curation.

To assess the quality of assemblies, we rely on the movements of k-mers^1^ across the taxonomic tree by addition of a given sequence to a Kraken2 database (Methods 4.5). The varying topologies of the families, however, add a bias to the notion of k-mer movement up the tree. A k-mer that moves one node up its branch within a family with a max depth of `6` is a relatively less damaging movement than the information lost for a node of max depth `1` moving the same amount. Sup. S2 illustrates a set of viruses, each reassigned to rank `S2`, with this kind of discordance: Influenza A (`11320`) reaches `3` levels with a subtree size `112,867`, compared to SARS-CoV-2 which is itself a leaf node. A deep family will tabulate more total k-mer distance movement than a shallow family which biases scoring. We therefore normalize raw tree distances by each family’s clade depth (Methods 4.4.1) to keep movement metrics comparable across families, regardless of how finely each clade is resolved in the taxonomy.

### 2.3. Quality metrics improve information gain and contamination control in databases

Through quality assessment of GenBank nucleotide viral accessions, we generate several databases for comparison (Methods 4.5). Database *S* is the standard database consisting solely of RefSeq genomes, *V* is RefSeq with 5,749,833 added genomes, while *A, B*, and *C* are the result of filtering and label reassignment of those 5.75M sequences.

The general goal of filtering was to strategically add reference genomes to the standard RefSeq database (*S*) such that we maximize k-mer information while minimizing perturbation to the original database. Fig. 2 shows how each database performs in this regard, by using each database to classify the GenBank viral sequences. Each sequence then has an LCA mapping from that database. Comparing each k-mer’s LCA to the sequence’s assigned label allows us to determine its expected position if it were added to the database (independent of other sequences). The standard database (*S*) lacks a significant number of k-mers in GenBank, which would lead to increased false negative results when classifying circulating strains in read samples, while *V* minimizes unseen k-mer rates but drastically increases the movement of k-mers up the tree, implying an increase in contamination from accessions added with poor quality. Database *A*, which includes leaf taxa with `0` added distance, finds a balance in the metrics between *V* and *S*. Databases *B* and *C* iterate on the construction of *A*; the new k-mer information allows for a reassessment of sequence quality, allowing for a further reduction in unclassified k-mer rates while maintaining, and even slightly improving, added distances per k-mer rates.

**Fig. 2.**
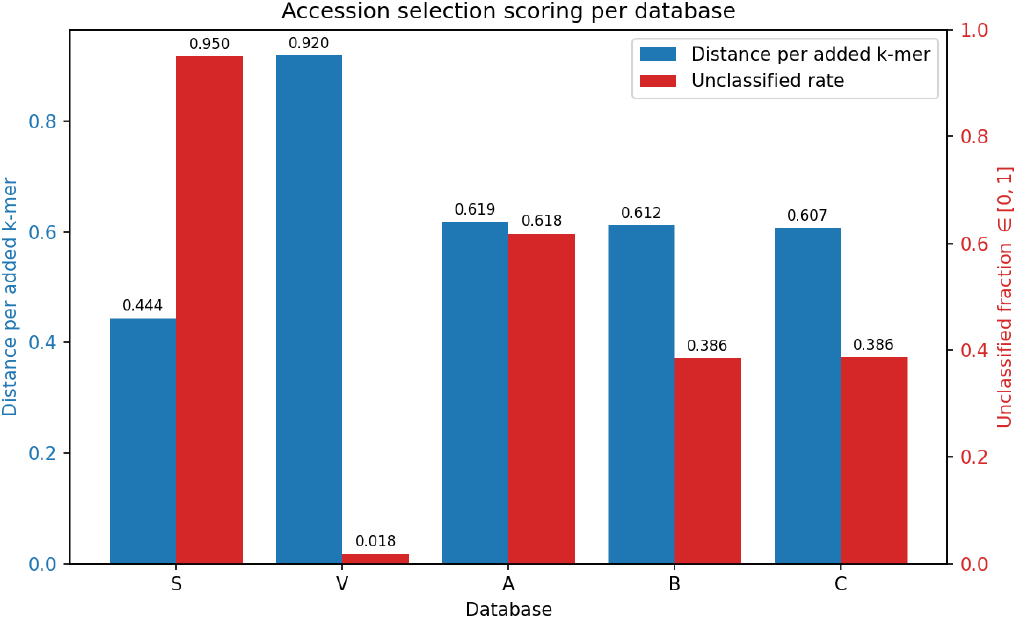
Classifying NT viruses with each database and using taxutils to score the perturbation allows for improvement in unclassified k-mer rates (red) while minimizing perturbation from each added k-mer (blue). An ideal database minimizes both metrics, assuming perfect quality in RefSeq.

For each database, Sup. S3 examines the contribution of different taxa to the distance added and how many unclassified k-mers remain for those taxa in each database because of their exclusion. Caudoviricetes (2832643) consistently appears as a top contributor of added distance. This makes intuitive sense: phage k-mers overlap with the bacteria they infect, so their addition pushes informative bacterial k-mers up to the root. Filtering them out, however, means we leave the unseen phage k-mers unclassified. A similar behavior occurs in HIV 1 (11676) with *Homo sapiens* (9606). Consistently low Monkeypox (10244) scoring leads to many of its accessions being left out of *A, B*, and *C*, raising the profile of its unclassification rate. Even in *V* we notice its high contribution which implies many k-mers were masked in the build and therefore excluded from the index; this implies a systematic issue of low quality regions in the Monkeypox reference genomes.

Using Kraken2 inspect, we determine the number of minimizers added to each taxon in the tree in each database. Fig. 3 illustrates the movement of minimizers away from leaf nodes in the standard as well as the number of newly seen leaf taxa with the added sequences. While *V* recovers only 51–67% more novel leaf taxa than *A, B*, and *C*, it displaces roughly `7.5×` more minimizers from their original leaf positions in the standard, indicating that its less selective filtering incurs a disproportionate cost in database perturbation relative to its gain in coverage.

**Fig. 3.**
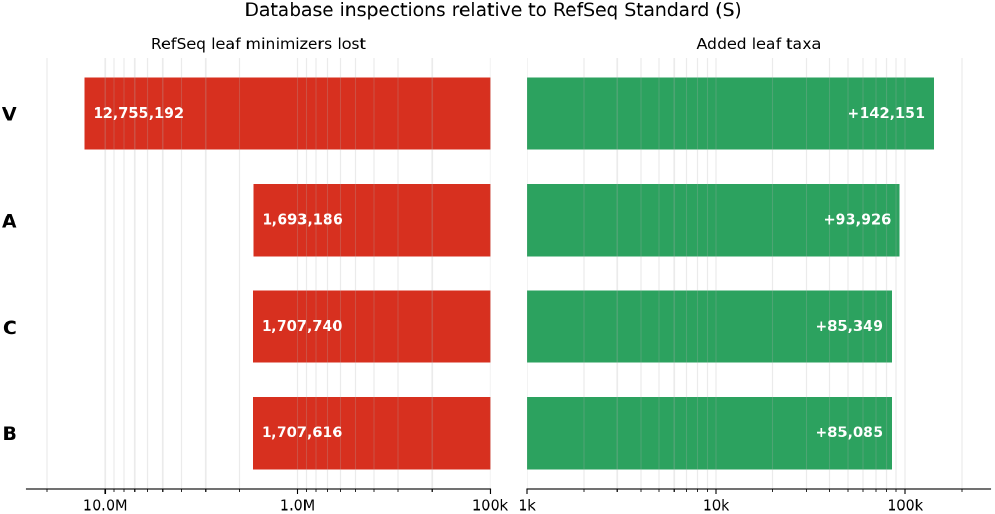
Adding sequences to a database involves a tradeoff between coverage of previously unseen taxa (green) and movement of minimizers away from their original position in the standard database (red).

## 3. Discussion

We have shown a complex analysis of taxonomic data and accessions performed through simple usage of `taxutils`. Tedious groundwork of taxonomic bookkeeping is simplified through trustworthy regex accession parsing, rank correction, tree traversal, topology summarization, and map construction. Using these operations, we diagnose rank incongruities, surface underrepresented viral families, and assemble sets of sequences to recover lost k-mers while preserving the trusted genomic information from the RefSeq curation.

The database results argue for a disciplined notion of quality. Adding all candidate sequences without filtration (*V*) minimizes unseen k-mers but buys that coverage at a disproportionate cost in perturbation. Databases *A, B*, and *C* navigate this tradeoff, and the reassessment enabled by each successive build lets *B* and *C* lower unclassified rates while minimizing perturbation up the LCA tree. We acknowledge that not all upward movement is contamination. An advantage of LCA mapping, beyond space conservation, is to keep generic k-mers from acting as unique identifiers. We therefore treat the standard RefSeq database as a reasonable ground truth, and acknowledge that some k-mer movement is healthy genomic information. Healthy but generic k-mer movement, however, exposes structural problems in the taxonomy, for instance with addition of phage k-mers (Caudoviricetes). These sequences in the viral subtree (10239) pull informative bacterial signal (2) to the root.

Label reassignment serves this same information gain: choosing a child label that minimizes a sequence’s k-mer movement up the tree reduces overall information loss. This selection is a heuristic to choosing subtype labels according to average nucleotide index. In the problem of accession selection, we hint at a min-max optimization problem. For the sake of keeping focus on illustration of `taxutils` utility, we avoid complicating the selection procedure and leave a more interesting optimization formulation to future work.

The intention of this work was to simplify the process of managing the necessary data for usage in common bioinformatics tools. The advent of coding agents, in particular, means users need just enough trustworthy base code for a model to assemble customized scripts around it. A codebase with verified function outputs can be relied on when an LLM calls them, minimizing outsourced decision-making to the LLM. `taxutils` was designed with this modern workflow in mind. Each of the example scripts in https://github.com/SwabSeq/taxutils/tree/main/examples was produced this way, using the provided SKILL.md. When the next tool demands its own idiosyncratic accession, taxon, or rank mapping, that is precisely the friction `taxutils` is meant to mitigate.

## 4. Methods

The `taxutils` functionality revolves around a self-named object; on construction, the relevant taxonomic data structures are loaded with access to various functions for operation on that data. We recommend binding the object to a convenient variable; throughout, we use `tu = taxutils(low_-memory=False)` and refer to it as `tu`.

### 4.1. Rank correction

To address the inconsistencies in taxonomic rankings, `taxutils` parses NCBI’s `nodes.dmp` into a DataFrame (accessed via `tu.nodes`) and augments it with several new columns, including `rank_code`, `rank_base`, `rank_idx`, and `new_rank`. Canonical ranks (`R`, `D`, `K`, `P`, `C`, `O`, `F`, `G`, `S`) are used as anchors whenever they fall deeper than their corrected parent rank. When a node instead carries a noncanonical label (e.g. `no rank` or `clade`), or a canonical label that is not deeper than its parent, it inherits its position from the tree: it is assigned a subrank one level below its parent’s code, e.g. `S2`, `S3`, or `F2`, denoting one and two levels below species (`S`) and one level below family (`F`), respectively, following the precedent of Kraken2 (Wood et al., 2019). The canonical name corresponding to the base of the corrected rank is stored in `new_rank`. Users can then call `tu.get_branch(taxon)` to trace a taxon’s lineage from root to taxon, `tu.higher_-than_rank(taxa, rank)` for a per-taxon boolean indicating whether each taxon sits above a given rank, and `tu.get_rank_order()` for the canonical rank ordering.

### 4.2. Calculating distance between nodes

Assuming the taxonomic tree has unit-length edges, the distance between two nodes *a* and *b* is measured through their lowest-common-ancestor `LCA(`*a, b*),

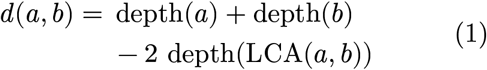

where `depth(`*v*) is the number of edges from the taxonomy root to node *v*. Equivalently, *d*(*a, b*) is the sum of the path lengths from *a* and *b* up to their common ancestor.

### 4.3. Measuring taxon clade topologies

Each of the following statistics can be calculated with `tu.topology(`*a*). To extract tree-structure details for an input taxon *a*, an anchor node *A*(*a*) is first selected according to a specified canonical rank, where we find the nearest ancestor with the selected rank, e.g., the closest family ancestor when `anchor_rank=“F”`,

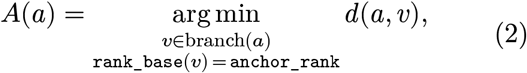

falling back to *a* itself if no such ancestor exists.

Then let *T* (*a*) be the subtree rooted at *A*(*a*) (the anchor and all descendants, as returned by `tu.get_subtree`). The depth of a descendant *x* below the anchor is

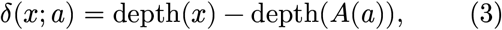

so δ(*A*(*a*); *a*) = 0. Write *N*(*a*) = |*T* (*a*)| for the clade size and ℒ(*a*) for the set of leaf taxa, those with `deg`^+^(*x*) = 0.

#### 4.3.1. Vertical structure

Several statistics describe the vertical extent of *T* (*a*): the clade size *N*(*a*), the terminal count |ℒ(*a*)|, and the maximum and mean descendant depth below the anchor,

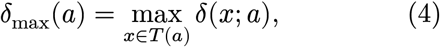

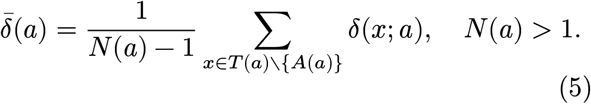

Together these distinguish compact clades 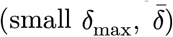 from deeply nested clades carrying many intermediate taxonomic levels.

#### 4.3.2. Branching structure

The broadest branching event is the maximum out-degree,

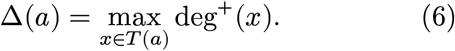

How frequently branching occurs is captured by the fraction of taxa with at least one descendant, equivalently one minus the leaf fraction,

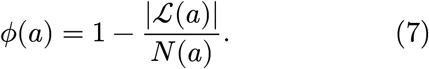

Finally, to characterize imbalance, let *c*_1_, …, *c*_*k*_ be the immediate children of the anchor, with ∑_*i*_ |*T* (*c*_*i*_)| = *N*(*a*) − 1. The fraction of descendant taxa in the largest immediate child branch is

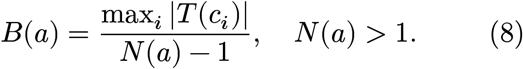

Values *B*(*a*) → 1 indicate diversity concentrated in a single branch, while smaller values (*B*(*a*) → 1/*k* when balanced) indicate a more evenly distributed topology.

### 4.4. Accession quality from LCA mapping

In the context of LCA k-mer databases, we ascribe the notion of quality to be the amount of perturbation induced by addition of new k-mers to some existing index. For a candidate sequence labeled *a*, we can lookup its current k-mer LCA mappings which we call ℬ, namely, the set of classified taxa observed among the sequence’s k-mers, excluding unclassified or masked k-mers. For each observed taxon *b ∈* ℬ, let *c*_*b*_ be the number of k-mers assigned to *b*, and let *l*_*b*_ = LCA(*a, b*) be the lowest common ancestor shared by the assigned label and the observed taxon. If we were to add this sequence to the current index, each k-mer’s LCA would move up the tree some distance, assuming *a* ≠ *b*. We can decompose each k-mer’s movement into two components. The *upward* distance is the ascent from the assigned label *a* to the lowest-common-ancestor *l*_*b*_,

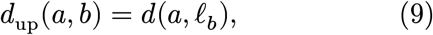

and measures how far one must climb out of the assigned branch before reaching the LCA. The *lateral* distance is the remaining descent from *l*_*b*_ down to the observed taxon *b*,

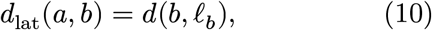

capturing divergence into a different lineage once the shared ancestor is passed. Their sum is the full tree distance between the two taxa,

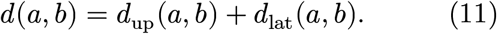

Across the contig, the total distance moved is the count-weighted sum over all classified k-mer taxa,

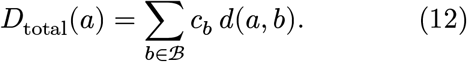

Letting 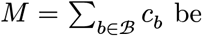 be the total number of classified k-mers, the^*b∈*^a^ℬ^verage distance moved is

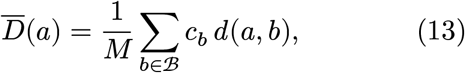

and the variance in per-k-mer movement is

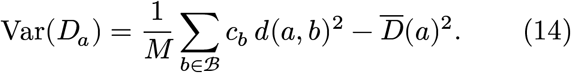

#### 4.4.1. Normalization of LCA movement

Because of varying topologies in different taxa clades, the distance a k-mer moves can be arbitrary when comparing between two distinct clades. We therefore introduce a scaling factor denoted *τ*_*a*_, defined as the 95th percentile of positive descendant depths (Eq. (3)) below the anchor:

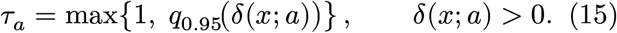

A distance of several edges may be large in a shallow family but typical in a deeply nested family, so we use *τ* to preserve tree-based movement information while reducing the influence of differences in clade depth and branching structure. Since *τ*_*a*_ depends only on *a*, it factors out of the sum. The total normalized distance moved by introducing a new label *a* is therefore

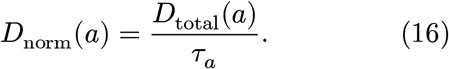

In the following experiment, we use the nearest family ancestor (`anchor_rank=“F”`) when calculating *τ*.

The notion of quality via perturbation, given each of these metrics, can take a variety of interpretations. One notion of quality is the amount of normalized distance that a sequence adds to the index. A high total normalized distance corresponds to high perturbation to the existing k-mer index, while a total of 0 corresponds to no perturbation.

An ideal sequence is one that adds close to 0 distance but contributes new information (previously unseen k-mers) to the index. Some added distance is inevitable, so we also rely on the variance which answers the amount of “purity” in the movement. That is, whether the added k-mers are moving to the same or different nodes. Finally, we care more about movement up the branch, rather than any movement down to a cousin node in the same clade, so in tie-breaking cases we rely on upward and exclude lateral distance.

### 4.5. Database construction with quality metrics and label reassignment

Using the described metrics in the previous section, we strategically build databases to minimize the amount of perturbation we introduce to a new database on top of a trusted baseline. We consider a candidate set of 5,749,833 viral sequences from NCBI’s nucleotide database and perform filtering procedures using our metrics. The selection of these candidates is described in Methods 4.8. In addition to the assigned labels for each sequence from NCBI, we can enumerate new labels using the LCA mapping from the previously constructed index and calculate distance metrics with new candidate labels. The assigned taxon label for that accession would be the one that minimizes the selected distance metrics, prioritized as described in Methods 4.4.1.

#### 4.5.1. Database construction procedure

To illustrate the advantage of this approach, we construct five databases. We first build an updated standard database (*S*) using a recent release of RefSeq (Methods 4.8). We then expand the standard database (*S*) by adding all 5.75M viral sequences without filtration to create a database *V*.

For the next database, we classify the 5.75M candidates with database *S* to get LCA mappings and calculate quality metrics with `taxutils`. We then filter for sequences labeled at leaf taxa with 0 added distance to the tree. The remaining 1,025,470 sequences from the original 5.75M are used for another expansion of *S*, denoted database *A*. This database ensures that the lowest branches of the tree are filled while the original tree remains unperturbed.

A fourth database, denoted *B*, is constructed by using LCA mappings from *A* on the candidate sequences. For each candidate we require at least one classified k-mer (`total_kmers > 0`) and retain only sequences whose quality metrics fall below the upper outlier threshold, defined per metric as *Q*_3_ + 1.5 (*Q*_3_ − *Q*_1_), where *Q*_1_ and *Q*_3_ are the first and third quartiles. We apply this criterion jointly to the total normalized distance, the average normalized distance, and the distance variance, discarding any sequence that is a high outlier in any of the three. The sequences passing all filters form the expansion used to construct database *B*.

Finally, a database *C* is constructed using label reassignment. With the information added from database *A*, previously unseen k-mers in the standard database are now visible. The quality scores are based on NCBI assigned taxon labels, but it is possible that some existing LCA mapping from database *A* proposes a label that includes useful k-mer information while preserving more information from the RefSeq curation than the original NCBI label. We therefore propose enumerating each possible label that appears in the LCA mapping for each accession and assessing the quality score given each new label. For each taxon hit in the LCA mapping of the candidates from *A*, we create a new candidate label for that sequence. We then assign new labels with the criteria that (1) the new label is within the subtree of the original label and (2) the new label has the minimum normalized added distance across all sequence candidates. The result is 280,604 relabeled sequences of the total candidates. With these new mappings, we perform filtering identical to that in *B* but with a new `seqid2taxid.map` file for the Kraken2 build.

#### 4.5.2. Quality scoring

We measure performance by the amount of contamination and information added. For contamination, we calculate how many k-mers from the original RefSeq viral library are moved off of their original mapping, which we count as detrimental perturbation. To measure how much information is added, we count the number of k-mers left unclassified over the entire NT viral FASTA. A high-quality, high-information database leaves fewer k-mers unseen while preserving as much of the original curation as possible. This leads to a tradeoff between preservation of the original curation and growth of information.

### 4.6. taxutils usage

The `taxutils` package is accessed through a single utility object. After importing the constructor with `from taxutils import taxutils`, a working object is created as `tu = taxutils()`. The constructor accepts arguments `accessions`, `low_memory`, `targets_json`, `refresh`, `wgs`, and `canonical`. If `accessions` is provided, accession-to-taxon mappings are loaded during initialization. By default, `low_memory=True`, so accession lookups are performed by scanning the compressed NCBI accession-to-taxon file directly; setting `low_memory=False` builds or reuses a local SQLite database, which requires more disk space and a slower first setup but is faster for repeated queries. The `targets_json` argument allows a custom pathogen target list to be supplied, but will be downloaded automatically by default. `refresh=True` forces the managed taxonomy, target, and accession resources to be redownloaded and regenerated for updates to the most recent NCBI data. The `canonical` argument is True by default, but if False, allows `taxutils` to overwrite non-specific Influenza A accession-taxa assignments with subtype specific mappings parsed from NCBI metadata.

The resulting `tu` object provides methods for accession parsing, accession–taxon mapping, tree traversal, topology summarization, and rank-aware filtering. Accession strings or FASTA-style headers can be standardized with `tu.parse_accession`, and accession-to-taxon mappings can be loaded with `tu.load_a2t`, to avoid unnecessarily loading the entire map into random access memory. The inverse operation, `tu.get_t2a(taxa, low_-memory=None)`, returns accessions assigned to one or more taxa. Taxonomic relationships are queried with `tu.get_branch(taxon)` for root-to-taxon lineage; `tu.get_subtree(taxon)` to return the taxon and its descendants; `tu.get_lca(taxon_-a, taxon_b)` returns the lowest-common-ancestor of two taxa. Accession parsing is crucial, and a robust regex construct is leveraged to ensure no records are lost when operating on FASTA files.

Taxutils provides five core functions for operating from the command line: `clean` finds replaces FASTA headers with the accession within, `deduplicate` removes duplicate accession entries in a FASTA, `extract` exports accessions found in FASTA headers, `filter` removes or keeps specified taxa using accession-to-taxon search, and `grep` extracts FASTA records matching requested accessions. A thorough explanation of additional functions and features in `taxutils` can be found in the README.md on GitHub.

## 4.7. Code availability

The software is available at github.com/Swab-Seq/taxutils under the MIT license. `taxutils` can be installed via PyPI (https://pypi.org/project/taxutils/) and conda from the bioconda channel (https://anaconda.org/bioconda/taxutils).

Installation instructions and usage examples are provided in the GitHub repository, including the analysis performed in this paper.

### 4.8. Data availability

The data and analyses presented in this paper are derived from publicly available NCBI resources. The `taxutils` package retrieves taxonomic data from https://ftp.ncbi.nih.gov/pub/taxonomy/taxdump.tar.gz and accession-to-taxon mappings from https://ftp.ncbi.nlm.nih.gov/pub/taxonomy/accession2taxid/nucl_gb.accession2taxid.gz. For the database analysis, for each of the four RefSeq divisions `fungi`, `bacteria`, `archaea`, and `viral`, we fetch genomes from https://ftp.ncbi.nlm.nih.gov/genomes/refseq/<division>/assembly_summary.txt, retaining only genomes with assembly levels of “Complete Genome” or “Chromosome”. The *Homo sapiens* reference genome is obtained from https://ftp.ncbi.nlm.nih.gov/genomes/refseq/vertebrate_mammalian/Homo_sapiens/assembly_summary.txt. Plasmid genomes are fetched from https://ftp.ncbi.nlm.nih.gov/genomes/refseq/plasmid/*.genomic.fna. Viral nucleotide sequences were obtained from the NCBI FTP server at https://ftp.ncbi.nlm.nih.gov/genomes/Viruses/AllNucleotide/AllNucleotide.fa. SARS-CoV-2 (2697049) sequences are excluded from candidates for practical space purposes. The accessions used to build each database are posted at https://github.com/SwabSeq/taxutils/tree/main/accs. The date of download for each data element was June 27, 2026, which corresponds with RefSeq release 235.

## Supplementary S1: Family-level representation in the viral tree

In table S1, the thirty viral families with the highest log-ratio of total to RefSeq-represented taxa are reported, quantifying the representation gaps discussed in the main text. For each family we list its total taxon count within the viral (10239) subtree, the number of represented taxa and backing accessions from RefSeq, and the corresponding represented and unrepresented fractions of taxa.

**Table S1.**
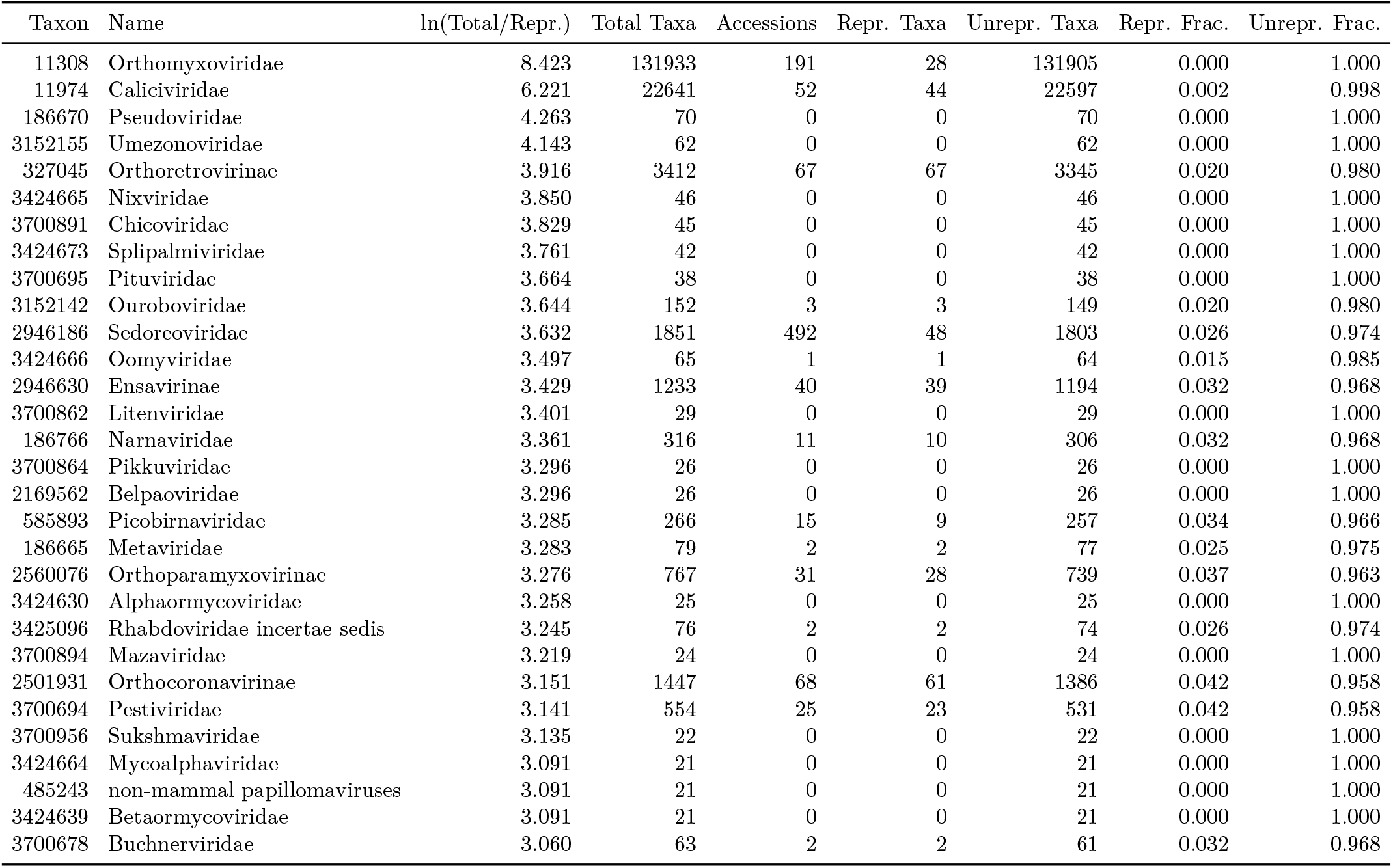
Family-level statistics in the viral RefSeq taxonomic tree. The log-ratio column uses a +1 pseudocount, ln((Total + 1)/(Repr. + 1)), so families with no represented taxa remain finite.

## Supplementary S2: Viral topologies

Although all eight respiratory viruses in table S2 share the reassigned rank `S2` (previously `no rank`), their taxonomic subtrees differ dramatically in size and branching structure, so statistics must be normalized before they are compared across taxa.

**Table S2.** Subtree size and shape vary by orders of magnitude from a single leaf (SARS-CoV-2) to 112,867 (Influenza A virus), and from flat, many-child polytomies to deeper balanced trees.

|  | SARS-CoV-2 | Rhinovirus A | Rhinovirus B | Rhinovirus C | Influenza A virus | Influenza B virus | Influenza C virus | Human RSV |
| --- | --- | --- | --- | --- | --- | --- | --- | --- |
| taxon | 2697049 | 147711 | 147712 | 463676 | 11320 | 11520 | 11552 | 11250 |
| original rank | no rank | no rank | no rank | no rank | no rank | no rank | no rank | no rank |
| rank code | S2 | S2 | S2 | S2 | S2 | S2 | S2 | S2 |
| n taxa | 1 | 87 | 40 | 63 | 112867 | 18580 | 312 | 49 |
| n leaves | 1 | 84 | 32 | 57 | 112713 | 18579 | 311 | 44 |
| max depth | 0 | 2 | 2 | 2 | 3 | 1 | 1 | 3 |
| mean depth | 0 | 1.011 | 1.175 | 1.063 | 1.999 | 1.000 | 0.997 | 1.918 |
| topology scale | 1 | 1 | 2 | 2 | 2 | 1 | 1 | 2 |
| max children | 0 | 84 | 31 | 57 | 36159 | 18579 | 311 | 25 |
| branching taxa fraction | 0 | 0.034 | 0.200 | 0.095 | 0.001 | 0.000 | 0.003 | 0.102 |
| top child fraction | 1 | 0.023 | 0.077 | 0.032 | 0.320 | 0.000 | 0.003 | 0.542 |

## Supplementary S3: Database comparison

**Fig. S1.**
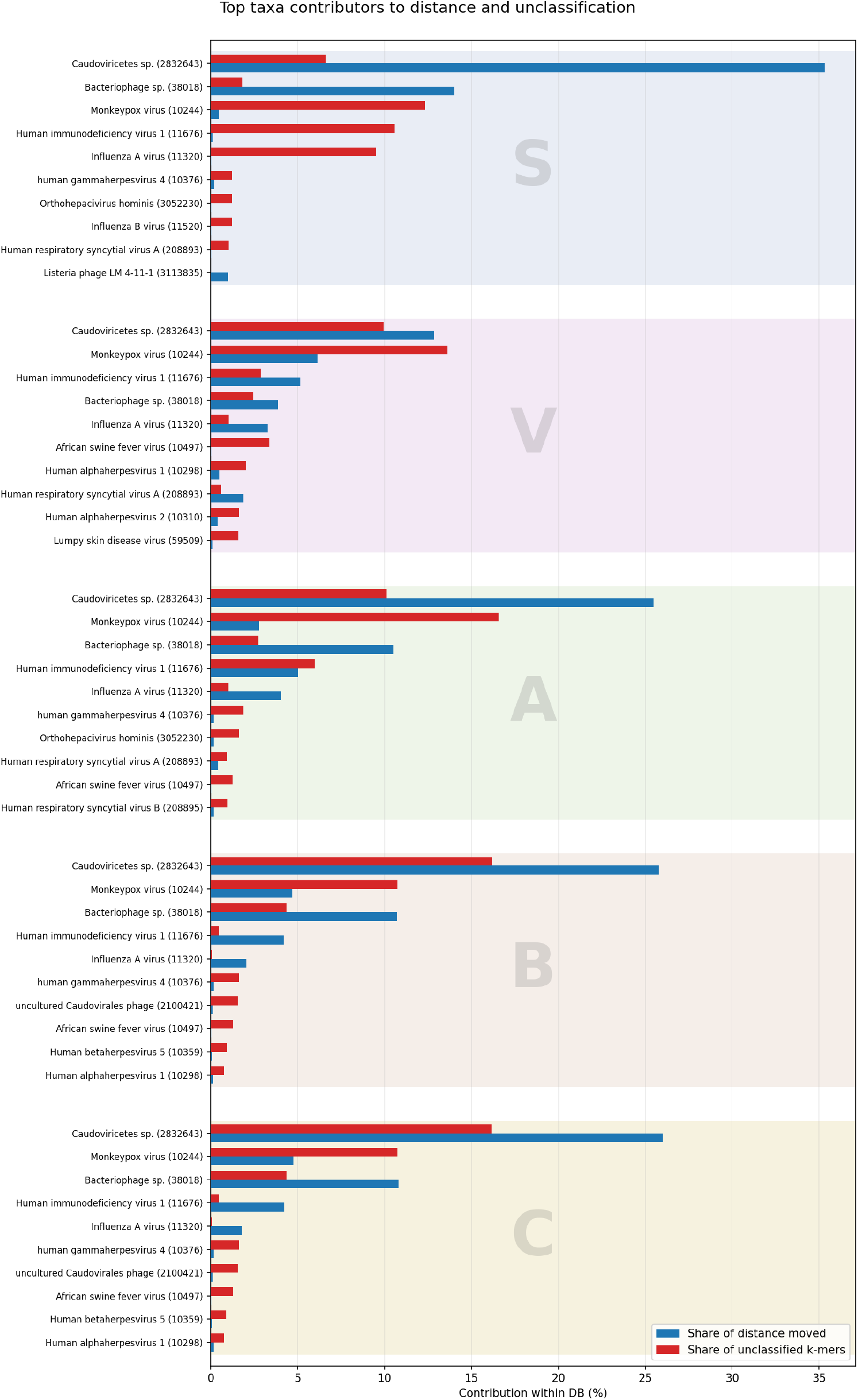
The relative total normalized distance and unclassified minimizer contribution across all accessions per taxon in each database. For example, sequences labeled 10244 (Monkeypox) contribute 6.12% of added normalized distance and 13.6% of the unclassified k-mers in database *V*.

## Footnotes

1 We use k-mer and minimizer interchangeably throughout.

